# PEPhub: a database, web interface, and API for editing, sharing, and validating biological sample metadata

**DOI:** 10.1101/2023.08.15.551388

**Authors:** Nathan J. LeRoy, Oleksandr Khoroshevskyi, Aaron O’Brien, Rafał Stepień, Alip Arslan, Nathan C. Sheffield

## Abstract

**Background:** As biological data increases, we need additional infrastructure to share it and promote interoperability. While major effort has been put into sharing data, relatively less emphasis is placed on sharing metadata. Yet, sharing metadata is also important, and in some ways has a wider scope than sharing data itself.

**Results:** Here, we present PEPhub, an approach to improve sharing and interoperability of biological metadata. PEPhub provides an API, natural language search, and user-friendly web-based sharing and editing of sample metadata tables. We used PEPhub to process more than 100,000 published biological research projects and index them with fast semantic natural language search. PEPhub thus provides a fast and user-friendly way to finding existing biological research data, or to share new data.

**Availability:** https://pephub.databio.org

## Background

The rapid pace of biological data generation has led to challenges with data sharing, storage, and integration [1–4]. Growing interest in data reusability and interoperability [5, 6] has led to new effort in improving biological data sharing and accessibility [7–9]. However, most effort has focused on biological *data*. Less emphasis has been placed on increasing the availability of biological *metadata* [10, 11].

As such, it is helpful to distinguish between *data* and *metadata*. In biology, *data* consists of experimental measurements or observations, while *metadata* describes the biological sample from which the measurements were derived. The sample metadata may include inherent, experimental, or analytical attributes about the sample. It might also describe the biology, treatments, experimental conditions, and data analysis parameters. Sharing complete biological *metadata* is important not only for integrated analysis, but also for discoverability [6]. There is a critical need for better tools and frameworks for sharing biological metadata.

To this end, tools and repositories have been developed to work with biological metadata [12–16]. However, they suffer from four main limitations: First, while metadata databases exist, they tend to focus on storage and retrieval [12]; none focus on simplifying user upload and editing of their own data. Second, metadata generally lacks a well-defined and supported structure. Previous methods tend to use a structure for a specific tool and data source [16–18], or leave the structure undefined altogether. Third, their search functionality is limited. Metadata search is generally limited to string matching or ontology searches. Finally, existing metadata services cannot easily be rebuilt and redeployed efficiently for custom use [19].

A recent advancement in biological metadata interoperability is Portable Encapsulated Projects (PEP), a framework that provides a standardized metadata structure, metadata validation, and programmatic metadata modifiers [20]. A PEP is a standardized sample table. The PEP framework provides a common infrastructure that links sample tables to analytical tools by removing the need for tedious and manual data preparation, mitigating the problem of metadata interoperability. However, there is no user-friendly web interface and API for sharing sample tables in the PEP ecosystem.

Here, we address these limitations with PEPhub: a database, web interface, search engine, and API for sharing, retrieving, and validating biological sample metadata. PEPhub provides several features that improve biological metadata interoperability, including: user- and machine-oriented interfaces, user editing and sharing, format conversion, metadata validation, natural language search, and containers for custom deployment. PEPhub advances the accessibility, discoverability, and reusability of biological sample metadata.

## Results and Discussion

### PEPhub instance and user interface

#### Public PEPhub instance

PEPhub is a web service for biological sample metadata. It is implemented as three major components that work together as modules: 1) a FastAPI web service; 2) a PostgreSQL database; 3) the PEPhubClient Python package, which provides Python and command-line interfaces to PEPhub (Figure 1A; Methods). To showcase the PEPhub software, we deployed a publicly available instance (see Availability). We used GEOfetch [21] to populate this public instance with over 150,000 projects (PEPs) derived from the Gene Expression Omnibus (GEO), with automated updates (Figure 1B, S1A; see Methods). PEPhub organizes projects by namespaces, corresponding to a user or organization on GitHub, thereby grouping related projects. PEPs are identified using a registry path in the form of `<namespace>/<project_name>:<tag>` (Figure 1C). The project name identifies a sample table. This naming convention allows convenient reference and versioning of sample metadata tables.

**Figure 1.**
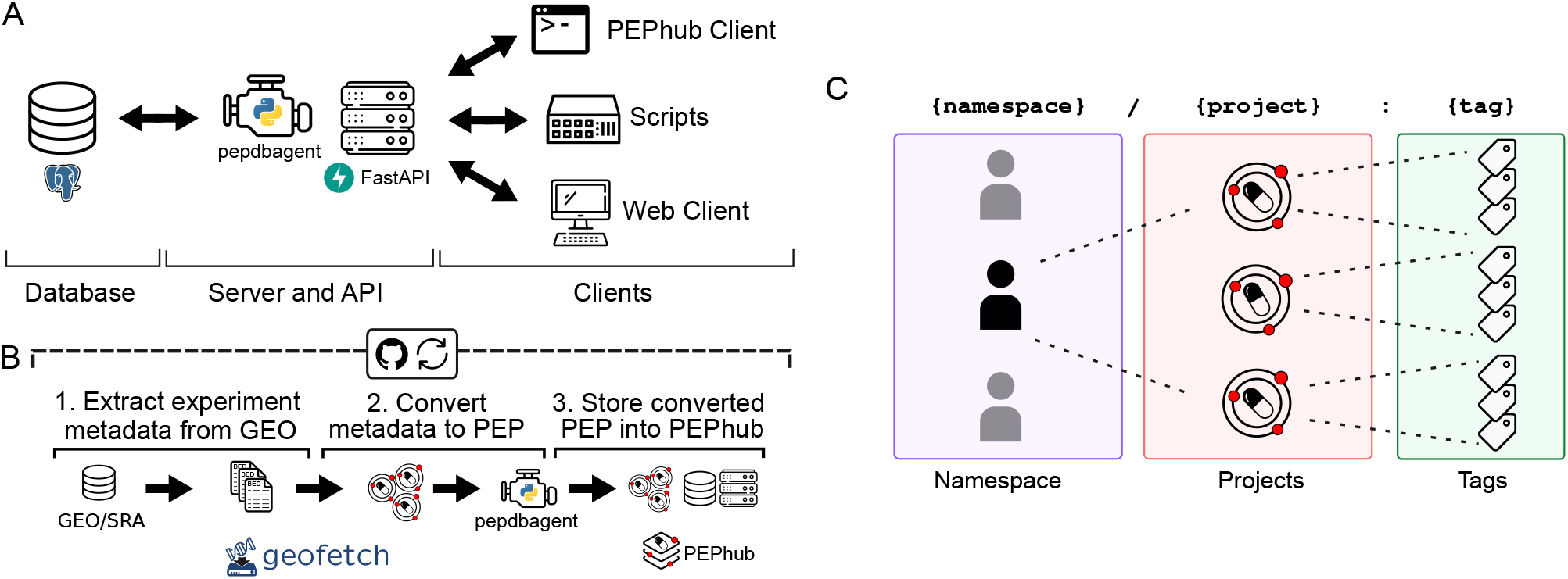
PEPhub high-level architecture and project identification strategy. **A**. PEPhub is backed by a Postgres database (left). It interfaces with the PEPhub server through a companion package called `pepdbagent` (middle). Web requests made by the web client or command-line interface are made via HTTP (right). **B**. Workflow for automated GEO-to-PEPhub transfer using GEOfetch. We take advantage of scheduled Github Actions to automate new discovery of GEO accessions to upload. **C**. PEPhub employs a `{namespace} / {project}:{tag}` nomenclature for sample table identification. Namespaces contain projects, which can be further distinguished with tags.

#### User- and machine-oriented interfaces

There are two primary interfaces by which users may interact with a PEPhub instance. First, the web interface provides access to PEP metadata for human browsing. It encourages data exploration and collaboration, making it easier for researchers to browse, search, submit, and edit PEPs. Second, the programmatic API allows other programs and scripts to interact with the server through HTTP requests. The API emphasizes the modularity of the PEPhub architecture and promotes interoperability with external software and services.

#### Format conversion

PEPhub provides programmatic interfaces to convert metadata into multiple formats. The standard PEP structure includes project-level attributes, a sample table, and a subsample table, which allows users to encode sample attributes with multiple values, such as sequencing reads with multiple file paths. By default, PEPhub offers the ability to convert this metadata into JSON, YAML, CSV, and plain-text formats (Figure 2A). To achieve this, PEPhub takes advantage of `eido`, a metadata validation engine written in Python [20]. Metadata conversion increases the interoperability of metadata, allowing it to fit into any analysis pipeline. Further, `eido` lets you write your own conversion functions, expanding the capabilities of a custom PEPhub deployment.

**Figure 2.**
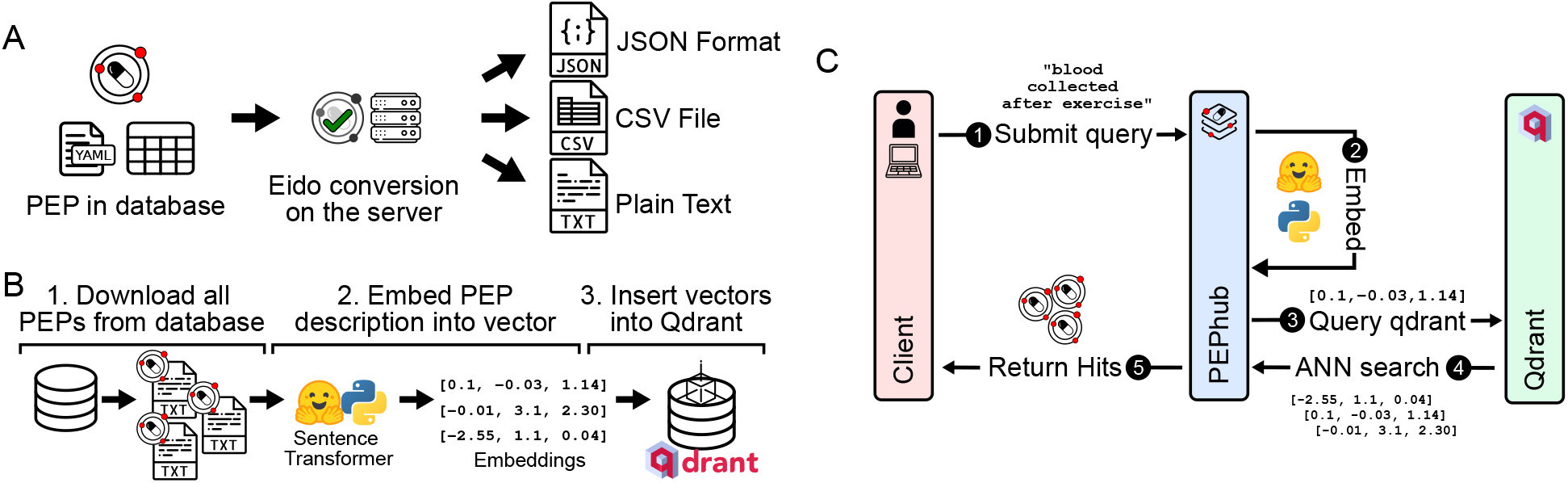
Metadata sharing, discovery, and accessibility features. **A**. PEPhub can convert metadata into `JSON`, `csv`, and `txt` output. **B**. Using a pre-trained sentence transformer, we periodically compute lowdimensional embeddings of all PEPs in PEPhub by mining text descriptions from the metadata. The resulting embeddings are then stored in Qdrant: a vector similarity engine and vector database. These embeddings are then compared against user-submitted queries. **C**. Searching for a PEP in pephub using vector search happens in five steps. First, the user submits a natural language query. Second, this query is embedded in real-time on the server. Third, the resultant vector is used to query Qdrant for nearest neighbors. Fourth, Qdrant responds with the most similar vectors it has stored. Finally, the hits are returned to the client submitting the query.

#### Natural language search

To improve biological metadata discovery, PEPhub provides a powerful natural-language search engine. The search engine is powered by pre-trained sentence transformers and a Qdrant vector database (Methods). We first use a sentence transformer to create lowdimensional vector representations of each PEP from the project-level and sample-level metadata attributes and descriptions. We store the resulting vectors inside a Qdrant vector database instance (Figure 2B). When a user provides a natural language search query, PEPhub transforms the query using the same sentence transformer in real-time, then queries the Qdrant API to retrieve the most semantically similar PEP vectors. Qdrant identifies similar PEPs by calculating nearest neighbors in vector space. PEPhub then returns the results to the client with their associated description and registry path (Figure 2C). PEPhub’s search engine uses a *semantic* approach, which provides several advantages:

First, the system returns results with similar meaning whether or not they include the terms of the original query. Second, it is tolerant of misspellings and is not limited to any ontology or taxonomy. Finally, because each PEP is represented as a vector, we can use highspeed nearest-neighbor algorithms to identify relevant PEPs, making the search very fast [22]. This method scales to millions of PEPs, and the speed is limited only by network speeds. Users may also tune results with limits, offsets, and relevance score cutoffs (Methods). To demonstrate the value of PEPhub’s semantic search, we show how some possible search terms like “childhood blood cancer” are able to retrieve more specialized related datasets (Figure S1B Supplementary Table 1).

#### Private and collaborative metadata upload and editing

While the natural language search and API access to standard structured metadata from GEO is valuable, one of the most important features of PEPhub is the ability for users to submit and edit their own PEPs. Users can submit and then edit their own PEPs on PEPhub through the API or through the web interface. To facilitate this, PEPhub also provides a robust authentication system. Users authenticate with PEPhub using GitHub, which provides user and organization namespaces. Users have read access to all namespaces but write access only to their namespaces. For PEPs with write access, users may mark them as *private* to restrict read access to only users with write access (Figure 3A). For example, **user1234** can edit all PEPs in the **user1234/** namespace. They may also edit all PEPs in the **org1234** namespace if they are a public member on GitHub. This ensures that only authorized users can access and modify private PEPs. By integrating authentication and authorization features, PEPhub provides a secure and controlled environment for users to interact with and manage their own PEPs while also facilitating the sharing and discovery of public PEPs to support collaborative research efforts.

**Figure 3.**
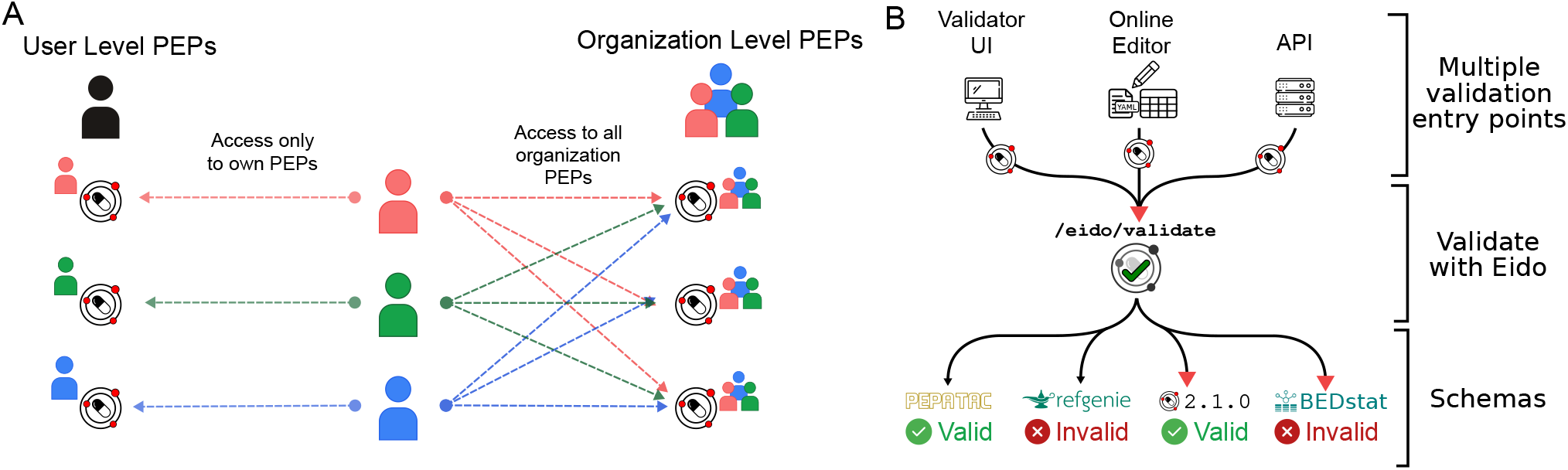
Metadata privacy and validation features. **A**. Users have read access to all namespaces but write access only to their namespaces (left). Other users are not permitted to modify a PEP in any user namespace other than their own. PEPhub implements organizations through GitHub. Members of an organization are automatically granted write access to all PEPs that belong to that organization (right). **B**. Validation on PEPhub is made easy with the integration of `eido`. PEPs in PEPub can be validated using either the web-based validator UI, the metadata builder, or programmatic endpoints.

**Figure 4.**
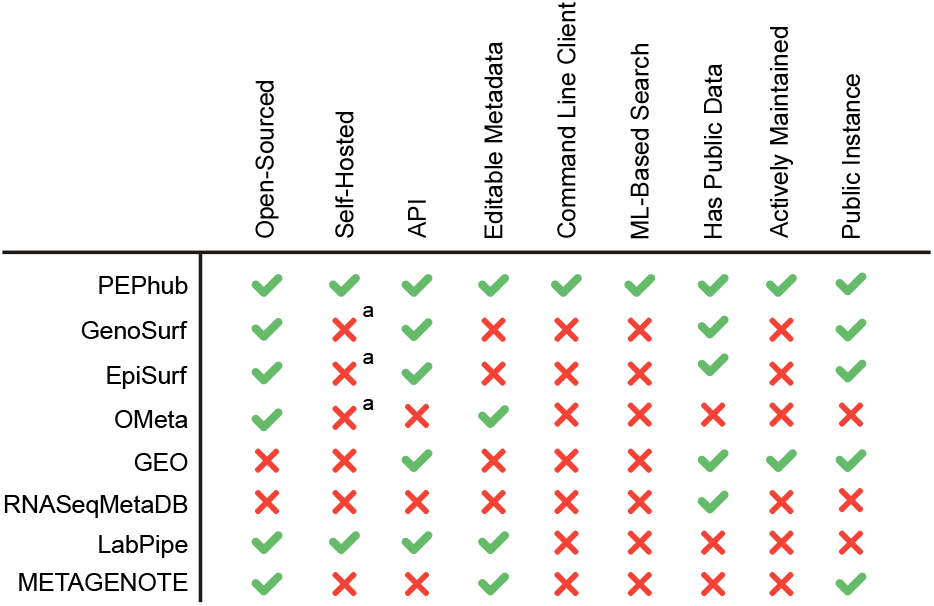
Metadata management comparison chart. PEPhub compares favorably to alternative metadata management systems. *a: While open-source, no clear documentation exists for self hosting an instance.

#### Metadata validation

PEPhub also provides metadata validation. We use `eido`, a PEP validation tool based on `jsonschema`, to validate on the server [20]. There are three ways to validate metadata through PEPhub. First, you may use the web-validator UI built with the server. With the web-validator UI, you may upload your own PEP or use PEPs stored on PEPhub and validate them against either PEPhub schemas or custom schemas. Custom schemas can be uploaded or pasted directly on the UI. Second, you can take advantage of the built-in metadata builder. When editing your PEPs, PEPhub validates the PEP after each save. The interface will propagate any errors to the user. Finally, there are validation endpoints at `/eido/validate`. These allow programmatic validation of PEPs (Figure 3B).

#### Comparison to other tools

Currently, several biological metadata management solutions exist to help alleviate the issue of metadata accessibility and interoperability; however, these solutions suffer from one or more limitations. One example is OMeta [19]. While it shares some features with PEPhub, it is not actively maintained, it lacks a public instance, and it lacks any documentation to start a private instance. Another example is RNASeqMetaDB [23]. Like PEPhub, RNASeqMetaDB aims to solve the problem of disorganized and limited access to sample metadata that are often published alongside the data itself; however, it lacks a currently running public instance and the source code is not available to reproduce the results. Finally, EpiSurf [18] and GenoSurf [12] are comprehensive genomics metadata search servers. However, they are limited in three ways: first, they don’t permit users to submit their own metadata and the database doesn’t appear to be regularly updated. Second, the software is not easily deployable. Third, their search system is based solely on biological ontologies, limiting the search space and flexibility of the search system.

PEPhub has several features that make it unique: First, it prioritizes allowing users to edit sample metadata stored on the server. This critical design decision positions PEPhub as not just as a place to *find* metadata, but as a place to manage and share your own. Second, it provides a full database, web API, and user interface for metadata management. Third, it provides the only metadata search engine that takes advantage of pre-trained sentence transformers for a powerful semantic search system. Fourth, it is the only tool that is open-sourced, automatically updated, actively maintained, and provides clear instructions to deploy a private instance. Together, these features make PEPhub a unique, flexible, tool that promotes the accessibility, findability, and interoperability of biological sample metadata.

PEPhub may also be compared to a Laboratory Information Management System (LIMS), a broad term with several interpretations. One difference is that LIMS tend to target management challenges related to physical sample handling in a wet lab, such as connecting to machinery, ordering reagents, and tracking physical samples through an experimental protocol. PEPhub could fulfill some of these functions, but in general, PEPhub targets a later phase of the experimental process: after data is generated, and the samples need to be analyzed and shared. PEPhub’s strengths are the ability to share data easily, in a more universal form, a public API, standardization, and searching. Thus, in many cases, it may make sense for PEPhub to live alongside a LIMS.

### Future development of PEPhub

We have several plans for PEPhub development. First, with some basic adapters, PEPhub could simplify the process of submitting data to public repositories, such as SRA or GEO. We are interested in working with interested parties to explore how to simplify the data submission process. Second, we plan to extend PEPhub to serve as a source for data analysis. We already have adopted our pipeline engine, looper, to retrieve sample tables from PEPhub. We plan to extend this functionality to other pipeline engines, such as Snakemake. Finally, PEPhub has potential to serve as a pipeline management dashboard, wherein processes could send updates to the server as pipelines run. We are currently developing software called `pipestat`, which provides a standardized mechanism for pipelines to report results, and we are exploring ways to link PEPhub with running pipelines such that they can be monitored directly from the web interface.

## Methods: Implementation and deployment details

### FastAPI web service

The PEPhub server is built with FastAPI, a web framework optimized for speed and high-performance. FastAPI is specifically designed for developing APIs. We chose FastAPI for its automatic data validation capabilities, built-in API documentation, and because using Python allows us to interface with existing Python infrastructure for metadata management we developed previously [20]. The FastAPI application uses our companion package `pepdbagent` to interface with a Postgres database. The user interface is built using React.js and TypeScript, and is packaged with the server.

### pepdbagent companion package

To manage project creation, fetching, deletion and insertion into the database, we developed a companion package called `pepdbagent`. `pepdbagent` acts as a simple wrapper around the popular Python object-relational mapper (ORM) SQLAlchemy to provide a convenient API for managing projects in our database. Both PEPhub itself and all maintenance scripts use `pepdbagent` to manage the PEPs stored inside the database.

### PostgreSQL database

Postgres is well-suited for storing the structured and unstructured data found in PEPs because it excels at both relational and document storage. The PEPhub database is comprised of three tables. The first table, `projects` stores the PEPs metadata and PEPs configuration. The `projects` table consists of twelve columns to store data like the project timestamp, project id, and the project configuration as a JSON blob. The other two tables: `samples`, and `subsamples` store samples and subsamples that are linked with `project.id` to a specific project in the `project table`. We host the public PEPhub database instance on the Amazon Web Services Relational Database Service (AWS RDS).

### PEPhubClient Python and command-line interface

To facilitate command-line interaction and third-party tools using PEPhub, we have developed PEPhubClient. PEPhubClient is a command-line interface and Python API that leverages the machine-oriented interfaces of PEPhub. Namely, the public API. PEPhubClient makes it easy to push and pull PEPs to any PEPhub instance. The command-line interface supports authentication to make authorized requests. This includes working with private PEPs, downloading PEPs, editing PEPs, and submitting PEPs. The PEPhubClient CLI is implemented in Python using the `typer` library.

### Containers for custom deployment

To standardize deployment and promote interoperability, we’ve packaged the PEPhub server and database as docker containers. These containers are made available on dockerhub. This makes it easy to launch your own instance of PEPhub.

### Populating PEPhub with biological sample metadata from GEO

To populate PEPhub, we developed a pipeline to ingest sample metadata from GEO. Our pipeline uploads PEPs from GEO in two steps. First, it identifies experiments that were added or updated in certain period of time to the Gene Expression Omnibus[7] using the GEOfetch Python API [21]. Second, GEOfetch downloads, formats and produces PEPs from GEO experiments that are later uploaded to the PEPhub database. Our database now stores more than 150,000 high-throughput sample tables from the last 10 years from Gene Expression Omnibus. We developed a pipeline that uses GitHub `schedule` actions to automate the download, formatting, and upload or re-upload of new project releases on GEO. Moreover, the pipeline includes an automatic check for successful previous uploads, ensuring that all GEO projects are consistently updated on PEPhub without the need for manual intervention.

### Natural language search

To support the text mining and embedding pipeline, we developed a companion tool called `pepembed` that embeds a database of PEPs and inserts them into a Qdrant database instance. For each PEP in the database `pepembed` does three things: first, it *flattens* the `yaml` representation of the sample metdata to create a continuous string; second, it utilizes an embedding model (e.g. sentence transformer) to produce a lowdimensional vector representation of this text; finally, `pepembed` will insert this vector in a Qdrant instance along with that PEPs namespace, name, and tag. We leverage GitHub actions to run indexing tasks periodically to ensure that all PEPs stay properly indexed, even if their data change. `pepembed` is open-source and available on GitHub.

### Authentication and authorization

PEPhub supports two authentication flows: authorization code flow and device code flow. Both take advantage of GitHub’s OAuth services. In addition, both authentication flows require users to login with GitHub via a web browser, upon which a code is returned. This code is then exchanged for a JSON Web Token (JWT) via a `POST` request which can be used to make subsequently authorized requests. While very similar, both flows exist to make it as easy as possible to integrate third-party software with a PEPhub instance.

## Availability of supporting source code and requirements

Project name: PEPhub

Project home page: https://pephub.databio.org

Operating system: Platform independent

Programming language: Python

License: BSD-2

bio.tools ID: pephub

SciCrunch RRID: SCR_024892

## Supplemental figures

**Supplementary Figure S1.**
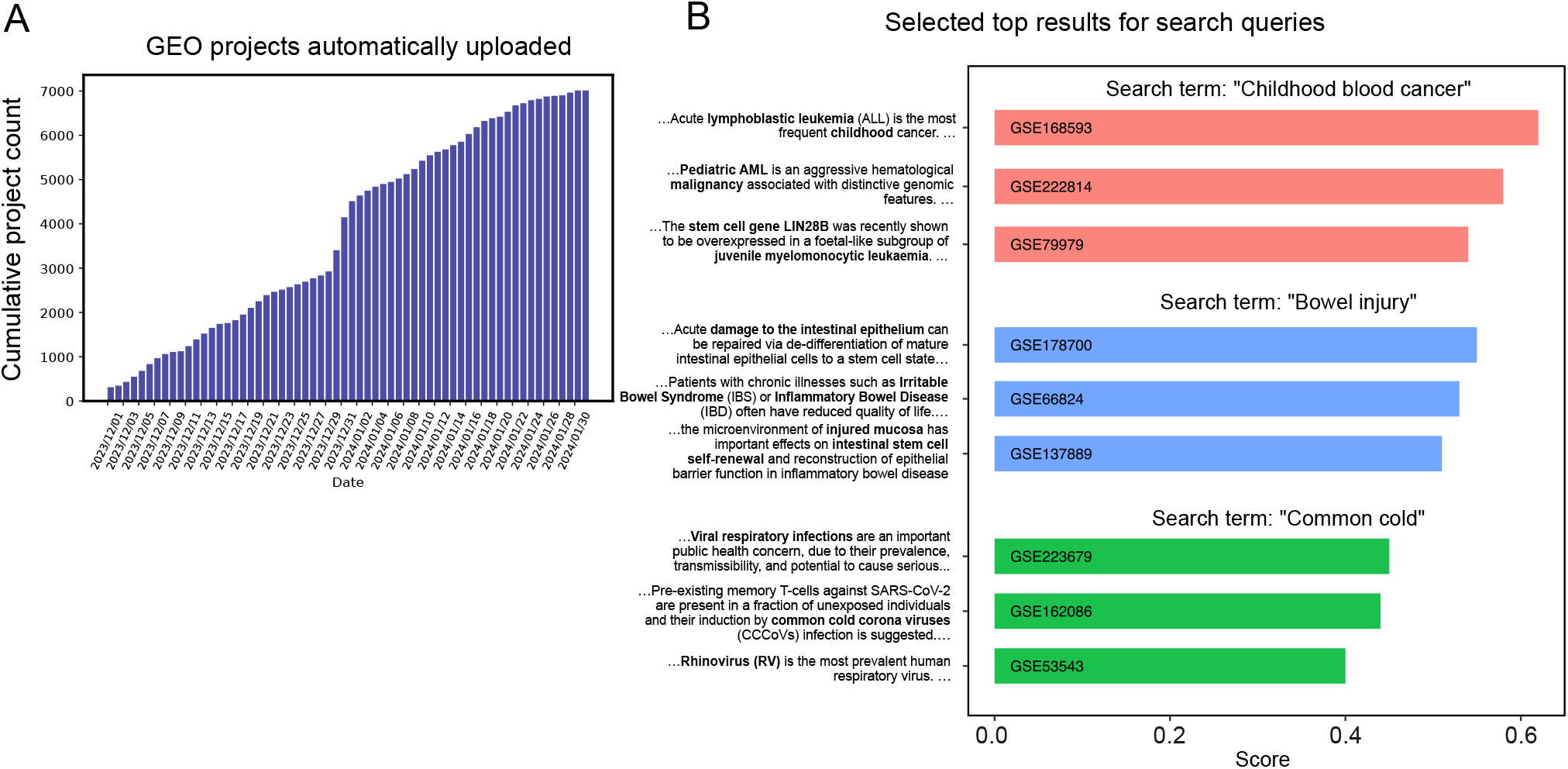
Analysis of PEP metadata and search. A) Barchart showing the cumulative number of new biological sample tables added to PEPhub from GEO, automatically. PEPhub automatically indexed more than 1000 new projects during this 3-week demo span. B) Illustrative search result scores selected from the top 10 responses returned by the PEPhub semantic search engine for biological search terms shown

## Supplemental text

### Search result examples

Illustrative search results from PEPhub. Search results from GEO and PEPhub for common queries. The PEPhub search engine returns results that are more relevant to the query than the GEO search engine. Moreover, the results returned by PEPhub are more diverse.

Search term: “Childhood blood cancer”

GEO results:

- … effects of MZ1 on multiple molecular subtypes of B-cell acute lymphoblastic leukemia cells (GSE217540)
- … PAF1 and FACT to drive high density enhancer interactions in leukemia (GSE202451)
- … PAF1 and FACT to drive high density enhancer interactions in leukemia… (GSE202450)
- … PAF1 and FACT to drive high density enhancer interactions in leukemia… (GSE202449)

PEPhub results:

- …genomic alterations in radiation-related breast cancer among childhood cancer survivors…(GSE62940)
- …analysis of CD10+/CD19+ pre B lymphoblasts from bone marrow and peripheral blood of B-ALL patients (GSE168593)
- …DNA methylation profiling predicts relapse in childhood B-cell acute lymphoblastic leukemia (GSE39141)
- …analysis of pediatric histiocytic sarcomas and anteceding hematologic malignancies (GSE109904)

Search term: “Bowel injury”

GEO results:

- tumors from mice fed diets excluding methionine/tryptophan/niacin (GSE246627)
- tumors from mice fed diets excluding methionine/tryptophan/niacin (GSE246626)
- Airway Microfold (M) Cells Emerge in the Post-IAV Lung (GSE244279)
- PEPhub results:
- Colonic mucosal injury responses (GSE164918)

Search term: “Common cold”

GEO results:

- ATAC-Seq of Batf-deficient pDC Transcriptomes…(GSE178410)
- Deterministic reprogramming of neutrophils in tumors (GSE244536)
- …Pancreatic Tumors Reveal Distinct Compartmentalisation of Neutrophil Subsets (GSE244534)
- …of neutrophil subsets in a mouse model of pancreatic cancer (GSE244531)

PEPhub results:

- A longitudinal study of natural respiratory viral infections (GSE223679)
- …temperature variation controls pre-mRNA processing and transcription of anti-viral genes (GSE193639)
- Influenzavirus serotype association to global whole blood transcriptional changes (GSE29385)
- The immune response and microbiota profiles during co-infection with P. vivax…(GSE144792)

## References

1. Volchenboum SL, Cox SM, Heath A, Resnick A, Cohn SL, Grossman R. Data Commons to Support Pediatric Cancer Research. American Society of Clinical Oncology Educational Book. 2017;746–52. doi:10.1200/EDBK175029.

2. Bui AAT, Van Horn JD. Envisioning the future of ‘big data’ biomedicine. Journal of Biomedical Informatics. 2017;69:115–7. doi:10.1016/j.jbi.2017.03.017.

3. Armit C, Tuli MA, Hunter CI. A Decade of GigaScience: GigaDB and the Open Data Movement. GigaScience. 2022;11:giac053. doi:10.1093/gigascience/giac053.

4. Xue B, Khoroshevskyi O, Gomez RA, Sheffield NC. Opportunities and challenges in sharing and reusing genomic interval data. Frontiers in Genetics. 2023;14. doi:10.3389/fgene.2023.1155809.

5. Wilkinson MD, Dumontier M, Aalbersberg IjJ, Appleton G, Axton M, Baak A, et al. The FAIR Guiding Principles for scientific data management and stewardship. Scientific Data. 2016;3:160018. doi:10.1038/sdata.2016.18.

6. Sheffield NC, Bonazzi VR, Bourne PE, Burdett T, Clark T, Grossman RL, et al. From biomedical cloud platforms to microservices: Next steps in FAIR data and analysis. Scientific Data. 2022;9:553. doi:10.1038/s41597-022-01619-5.

7. Edgar R, Domrachev M, Lash AE. Gene Expression Omnibus: NCBI gene expression and hybridization array data repository. Nucleic Acids Research. 2002;30:207–10.

8. Sloan CA, Chan ET, Davidson JM, Malladi VS, Strattan JS, Hitz BC, et al. ENCODE data at the ENCODE portal. Nucleic Acids Research. 2016;44:D726–32. doi:10.1093/nar/gkv1160.

9. Bourne PE, Bonazzi V, Dunn M, Green ED, Guyer M, Komatsoulis G, et al. The NIH Big Data to Knowledge (BD2K) initiative. Journal of the American Medical Informatics Association : JAMIA. 2015;22:1114. doi:10.1093/jamia/ocv136.

10. Leipzig J, Nuüst D, Hoyt CT, Ram K, Greenberg J. The role of metadata in reproducible computational research. Patterns. 2021;2:100322. doi:10.1016/j.patter.2021.100322.

11. Sheffield N, LeRoy N, Khoroshevskyi O. Challenges to sharing sample metadata in computational genomics. Frontiers in Genetics. 2023;14.

12. Canakoglu A, Bernasconi A, Colombo A, Masseroli M, Ceri S. GenoSurf: Metadata driven semantic search system for integrated genomic datasets. Database. 2019;2019:baz132. doi:10.1093/database/baz132.

13. Serna Garcia G, Leone M, Bernasconi A, Carman MJ. GeMI: Interactive interface for transformer-based Genomic Metadata Integration. Database. 2022;2022:baac036. doi:10.1093/database/baac036.

14. Masseroli M, Pinoli P, Venco F, Kaitoua A, Jalili V, Palluzzi F, et al. GenoMetric Query Language: A novel approach to large-scale genomic data management. Bioinformatics. 2015;31:1881–8. doi:10.1093/bioinformatics/btv048.

15. Davis S, Meltzer PS. GEOquery: A bridge between the Gene Expression Omnibus (GEO) and BioConductor. Bioinformatics. doi:10.1093/bioinformatics/btm254. 2007;23:1846–7.

16. Quin ñones M, Liou DT, Shyu C, Kim W, Vujkovic-Cvijin I, Belkaid Y, et al. “METAGENOTE: A simplified web platform for metadata annotation of genomic samples and streamlined submission to NCBI’s sequence read archive.” BMC Bioinformatics. 2020;21:378. doi:10.1186/s12859-020-03694-0.

17. Cappelli E, Cumbo F, Bernasconi A, Canakoglu A, Ceri S, Masseroli M, et al. OpenGDC: Unifying, Modeling, Integrating Cancer Genomic Data and Clinical Metadata. Applied Sciences. 2020;10:6367. doi:10.3390/app10186367.

18. Bernasconi A, Cilibrasi L, Al Khalaf R, Alfonsi T, Ceri S, Pinoli P, et al. EpiSurf: Metadata-driven search server for analyzing amino acid changes within epitopes of SARS-CoV-2 and other viral species. Database. 2021;2021:baab059. doi:10.1093/database/baab059.

19. Singh I, Kuscuoglu M, Harkins DM, Sutton G, Fouts DE, Nelson KE. OMeta: An ontology-based, data-driven metadata tracking system. BMC bioinformatics. 2019;20:8. doi:10.1186/s12859-018-2580-9.

20. Sheffield NC, Stolarczyk M, Reuter VP, Rendeiro AF. Linking big biomedical datasets to modular analysis with portable encapsulated projects. GigaScience. 2021;10. doi:10.1093/gigascience/giab077.

21. Khoroshevskyi O, LeRoy N, Reuter VP, Sheffield NC. GEOfetch: A command-line tool for downloading data and standardized metadata from GEO and SRA. Bioinformatics. 2023;btad069. doi:10.1093/bioinformatics/btad069.

22. Malkov YA, Yashunin DA. Efficient and robust approximate nearest neighbor search using Hierarchical Navigable Small World graphs. 2018. doi:10.48550/arXiv.1603.09320.

23. Guo Z, Tzvetkova B, Bassik JM, Bodziak T, Wojnar BM, Qiao W, et al. RNASeqMetaDB: A database and web server for navigating metadata of publicly available mouse RNA-Seq datasets. Bioinformatics (Oxford, England). 2015;31:4038–40. doi:10.1093/bioinformatics/btv503.

